# Expression status transition of *NOTCH1* accompanies chromatin remodeling in human early retinal progenitor cells

**DOI:** 10.1101/2023.07.10.548476

**Authors:** Yoshitoku Watabe, Sakurako Kobayashi, Takahiro Nakayama, Satoru Takahashi, Masaharu Yoshihara

## Abstract

**Objective:** The regulation of receptor expression is crucial for fine-tuned signal transduction. Notch signaling is a key signaling pathway involved in retinal development. Although the involvement of this signaling pathway in the differentiation of retinal ganglion cells has been documented, less is known about its involvement in earlier stages of retinal progenitor cell differentiation. To clarify the timing of Notch receptor expression in undifferentiated retinal progenitor cells and elucidate the possible involvement of chromatin remodeling in the regulation of Notch receptor expressions, we re-analyzed publicly available human fetal retina single-cell RNA-seq and ATAC-seq data.

**Results:** On days 59, 74, and 78, we observed *NOTCH1* mRNA expression in early retinal progenitor cells, which diminished at later stages of differentiation. Integration of single-cell RNA-seq and ATAC-seq revealed that chromatin remodeling in part of the *NOTCH1* locus was accompanied by transitions in its mRNA expression. Importantly, chromatin accessibility in the region upstream of *NOTCH1* depended on the differentiated cell types. These results suggest that chromatin remodeling may be involved in the differential expression of *NOTCH1*, although another type of Notch mRNA expression regulation may exist.

## Introduction

Signal transduction depends on the expression of receptors regulated at multiple levels. These regulations include chromatin remodeling and DNA-binding proteins, such as transcription factors and transcriptional repressors. Because fine-tuned signal transduction is necessary for development, it is important to clarify the regulatory mechanisms of receptor expression.

Notch signaling in mammals is dependent on the binding of five canonical DSL ligands and four Notch receptors [1]. It has been suggested that Notch loci are subject to chromatin remodeling under both normal [2-7] and pathological [8-16] conditions. In terms of developmental biology, the retina is a good model for investigating cellular differentiation involving Notch signaling [17]. The developed retina is composed of multiple cell types, including retinal ganglion cells, bipolar cells, photoreceptors, amacrine cells, horizontal cells, and Müllar glial cells, all of which originate from the retinal progenitor cells (RPCs). RPC is characterized by the expression of *Lhx2*, *Pax6*, *Rax*, and *Vsx2* while there are some additional marker genes for developing horizontal cells/retinal ganglion cells (*Onecut1*/*2*) [18], developing amacrine cells (*Elavl* genes) [19], developing photoreceptors/amacrine cells/Müllar glial cells (*Eef1a1*) [20], developing retinal ganglion cells (*Meis2*) [21], developing horizontal cells (*Lhx1*, *Ptf1a*) [22], and glial cells (*Pax2*) [23]. Of the four Notch receptors in mammals, Notch1, has been suggested to be involved in the maintenance of RPCs and differentiation into retinal ganglion cells [24]. However, it is unclear when and how Notch receptor expression is switched on and off in RPCs during early differentiation. Here, we re-analyzed a public multi-omics dataset of single-cell RNA-seq and single-cell ATAC-seq from three human fetal retinas [25] to clarify the timing of Notch receptor expression and examine the involvement of chromatin remodeling in this receptor expression switch.

## Methods

A single-cell multi-omics dataset (GSE183684) [25] was downloaded from the Gene Expression Omnibus (https://www.ncbi.nlm.nih.gov/geo/); data from days 59, 74, and 78 in the dataset were selectively used because they contained many undifferentiated retinal progenitor cells. The data were processed in Seurat version 5.1.0 [26] and Signac version 1.14.0 [27] pipelines in R version 4.4.1 on the Ubuntu 22.04.4 LTS environment. Pseudotime analysis and integration of the single-cell RNA-seq data and the single-cell ATAC-seq data was conducted by using the “FindTransferAnchors” function of Signac and Monocle3 version 1.3.7 [28], respectively. The conditions used in the analysis are provided in the GitHub repository.

## Results

### Re-analysis day 59 human fetal retina

First, early gestational stage samples were characterized (day 59). UMAP analysis of single-cell RNA-seq identified 13 clusters (Figure 1A), which were further characterized by marker gene expression (Figure 1B). This sample primarily contained RPCs with various differentiation statuses except for *PAX2*-expressing glial cells. Early RPCs (RPC1-3 and *MKI67*-expressing Proliferating RPC) were characterized by *LHX2*, *PAX6*, *RAX*, and *VSX2*. In addition, pseudotime analysis suggested that these RPCs differentiated into either *ONECUT1*/*2*-expressing, *ELAVL2*/*4*-expressing, or *EEF1A1*-expressing RPCs (Figure 1C). Next, we examined Notch mRNA expression and found that *NOTCH1*-*3* was expressed primarily in early RPCs, with *NOTCH1* and *NOTCH3* being the most prominent genes (Figure 1D). Then, we re-analyzed single-cell ATAC-seq data for this day 59 sample, which was mathematically integrated with its single-cell RNA-seq data by using the “FindTransferAnchors” function of Signac (Figure 1E). In the early RPCs (RPC1/2 and *MKI67*-expressing Proliferating RPC), we observed peaks between Chr9 136510000-136520000 of the *NOTCH1* gene, which diminished in the other RPC clusters. These results imply that Notch expression transition during RPC differentiation might be associated with the chromatin remodeling of *NOTCH1* (Figure 1F). However, we noted that chromatin accessibility of the upstream region of the *NOTCH1* locus remained high in *ONECUT*-expressing RPCs. In contrast to these populations, chromatin accessibility of the *NOTCH1* locus in *ELAVL2/4*-expressing RPCs was very low, which was consistent with the low mRNA expression in these RPC populations. In addition, the chromatin accessibility of the *NOTCH3* locus remained unchanged during RPC differentiation, although its mRNA expression diminished (Figure 1G). Since *NOTCH2* and *NOTCH4* were less prominent, the feature plots of the marker genes and the dot plots of the Notch genes from single-cell RNA-seq along with coverage plots of *NOTCH2* and *NOTCH4* from single-cell ATAC-seq are available in “Additional_file_1.”

**Figure 1.**
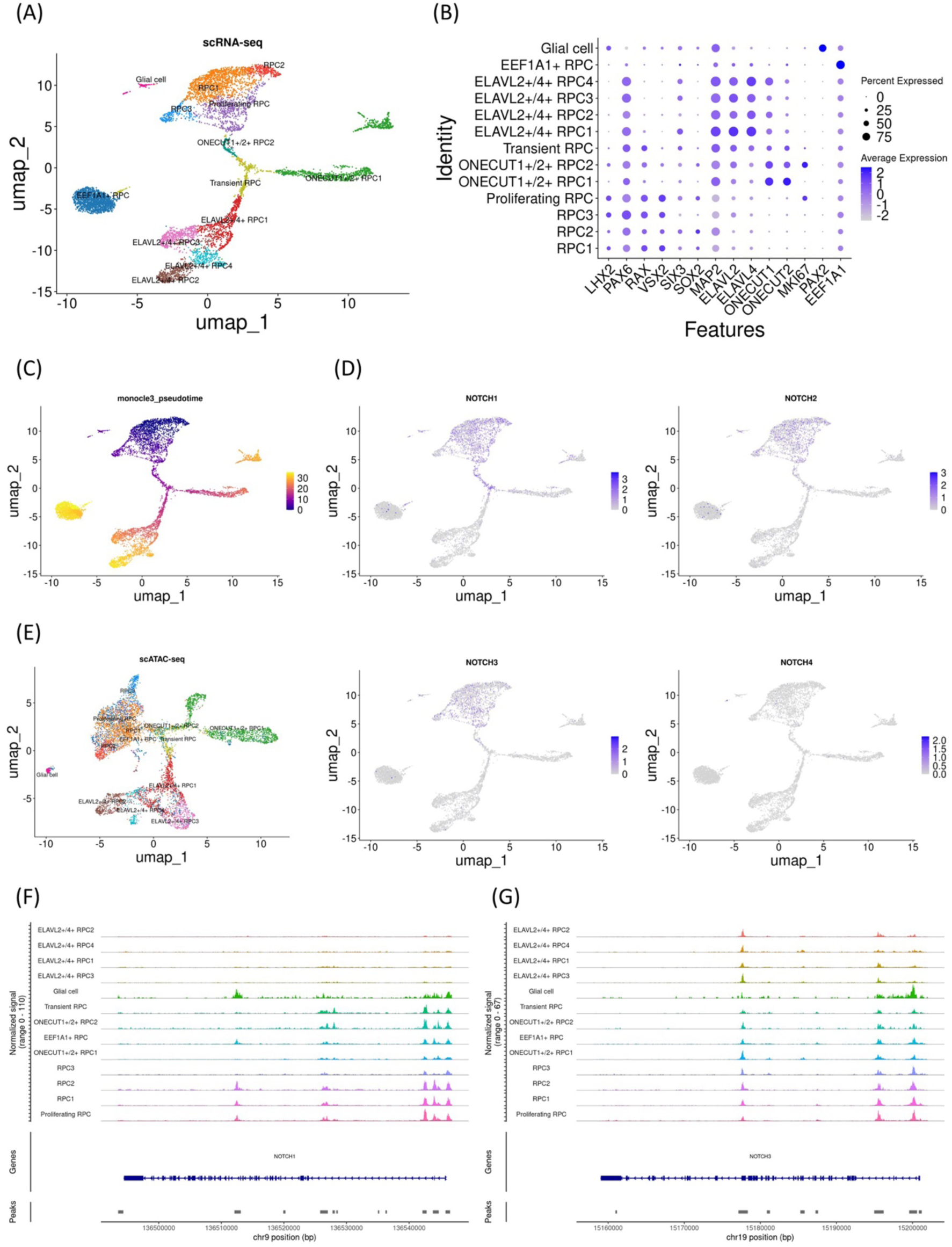
*NOTCH1* mRNA expression decrease and concomitant chromatin remodeling in the day 59 sample. (A) UMAP analysis of the single-cell RNA-seq data identified 13 clusters. (B) Dot plot of marker genes. Note that early RPC clusters expressed *LHX2*, *PAX6*, *RAX*, and *VSX2*. (C) Monocle3 pseudotime analysis for clarifying the differentiation status. (D) Feature plot of *NOTCH1*-*4*. Note that *NOTCH1* and *NOTCH3* expression was prominent in the early RPC clusters. (E) UMAP analysis of the single-cell ATAC-seq data which was integrated with the single-cell RNA-seq data. (F) Coverage plot of the *NOTCH1* locus showing the chromatin accessibility. Note that the upstream of the gene is on the right. (G) Coverage plot of the *NOTCH3* locus. Note that the upstream of the gene is on the right.

### Re-analysis day 74 human fetal retina

To examine whether the changes in Notch mRNA expression and chromatin accessibility of the Notch loci in the day 59 sample were representative, we investigated another early gestational stage sample (day 74). UMAP analysis identified 15 clusters (Figure 2A) that were further characterized by marker gene expression (Figure 2B). This sample primarily contained RPCs with various differentiation statuses. Early RPCs (RPC1-6) were characterized by *LHX2*, *PAX6*, *RAX*, and *VSX2*. In addition, pseudotime analysis suggested that these RPCs differentiated into either *ELAVL2*/*4*-expressing, *ONECUT1*/*MEIS2*-expressing, or *ONECUT2*-expressing RPCs (Figure 2C). *ONECUT1*/*MEIS2*-expressing RPCs differentiated into RPCs that markedly expressed *VSX2*. Next, we examined Notch mRNA expression and found that *NOTCH1*-*3* was expressed primarily in early RPCs, with *NOTCH1* and *NOTCH3* being the most prominent genes, similar to the day 59 sample (Figure 2D). We then re-analyzed the single-cell ATAC-seq data for this sample, which were integrated with the single-cell RNA-seq data (Figure 2E). In the early RPCs (RPC1-3), we observed peaks between Chr9 136510000-136520000 of the *NOTCH1* gene, which diminished in the other RPC clusters. These results confirmed that Notch expression transition during RPC differentiation might be associated with the chromatin remodeling of *NOTCH1* (Figure 2F). However, we noted that the chromatin accessibility of the upstream region of the *NOTCH1* locus remained high in *ONECUT*-expressing RPCs. In contrast to these populations, the chromatin accessibility of the *NOTCH1* locus in *ELAVL2/4*-expressing RPCs was very low, which was consistent with the low mRNA expression in these RPC populations. In addition, the chromatin accessibility of the *NOTCH3* locus remained unchanged during RPC differentiation (Figure 2G). In summary, similar to the day 59 sample, in the day 74 sample, a concomitant mRNA decrease and chromatin remodeling in *NOTCH1* was observed, while chromatin accessibility in the upstream region of the *NOTCH1* locus remained high in *ONECUT*-expressing RPCs. The feature plots of the marker genes and dot plots of the Notch genes from single-cell RNA-seq along with coverage plots of *NOTCH2* and *NOTCH4* from single-cell ATAC-seq are available in “Additional_file_2.”

**Figure 2.**
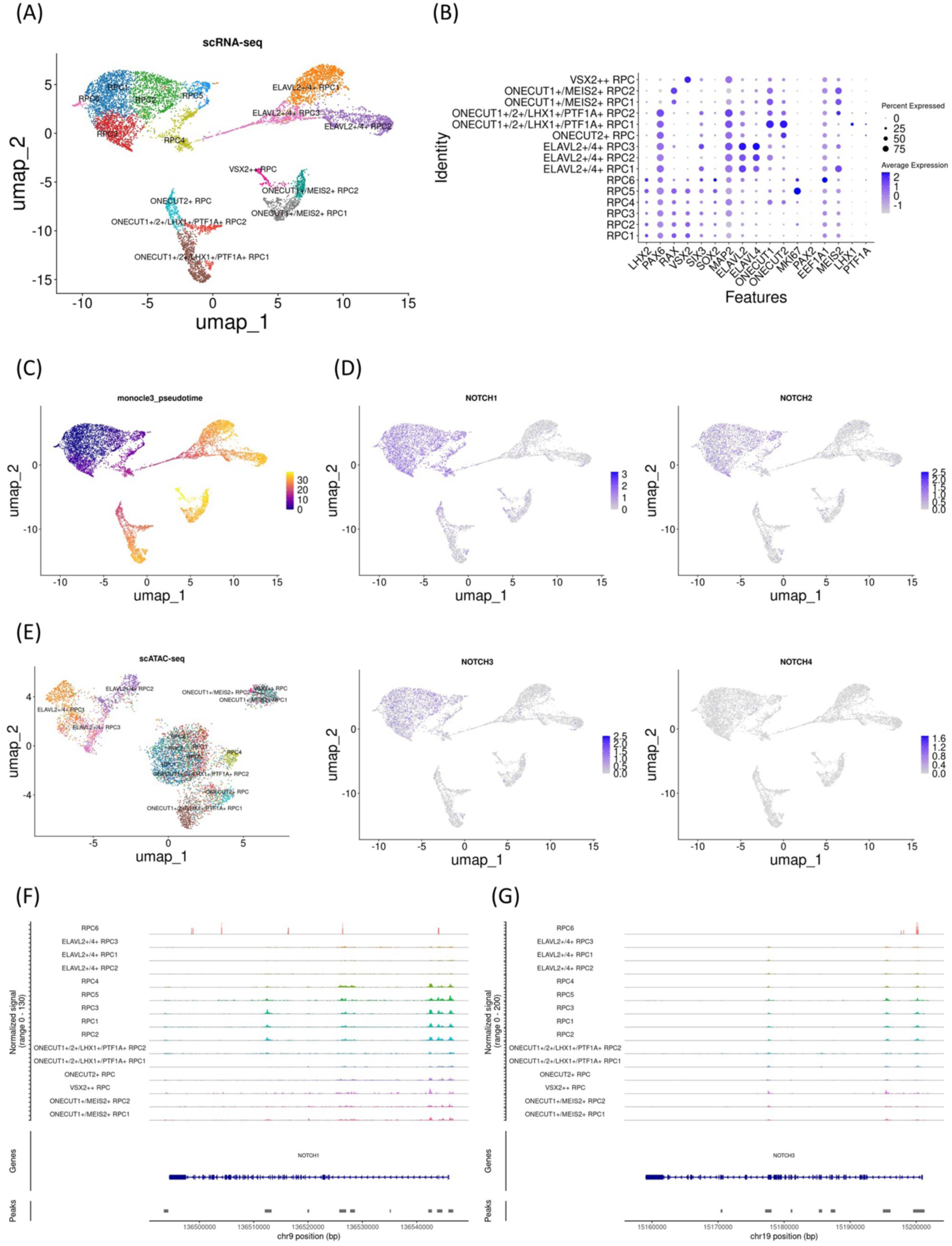
*NOTCH1* mRNA expression decrease and concomitant chromatin remodeling of the day 74 sample. (A) UMAP analysis of the single-cell RNA-seq data identified 15 clusters. (B) Dot plot of marker genes. Note that early RPC clusters expressed *LHX2*, *PAX6*, *RAX,* and *VSX2*. (C) Monocle3 pseudotime analysis. (D) Feature plot of *NOTCH1*-*4*. Note that *NOTCH1* and *NOTCH3* expression was prominent in the early RPC clusters. (E) UMAP analysis of the single-cell ATAC-seq data which was integrated with the single-cell RNA-seq data. (F) Coverage plot of the *NOTCH1* locus. (G) Coverage plot of the *NOTCH3* locus.

### Re-analysis day 78 human fetal retina

To further confirm the developmental changes in Notch mRNA expression and chromatin accessibility at the Notch loci, we investigated an additional early gestational stage sample (day 78). UMAP analysis identified 17 clusters (Figure 3A) that were further characterized by marker gene expression (Figure 3B). The sample primarily contained RPCs with various differentiation statuses. Early RPCs (RPC1-3 and *MKI67*-expressing Proliferating RPC1-2 cells) were characterized by *LHX2*, *PAX6*, *RAX*, and *VSX2*. In addition, pseudotime analysis suggested that these RPCs differentiated into *ONECUT1*/*2*-expressing RPCs, which later differentiated into *PTF1A* and *LHX1*-expressing RPCs, *MEIS2*-expressing RPCs, or *ELAVL2/4*-expressing RPCs (Figure 3C). Next, we examined Notch mRNA expression and found that *NOTCH1*-*3* was expressed primarily in early RPCs, with *NOTCH1* and *NOTCH3* being the most prominent genes (Figure 3D). We then re-analyzed the single-cell ATAC-seq data for the day 78 sample, which was integrated with the single-cell RNA-seq data (Figure 3E).

**Figure 3.**
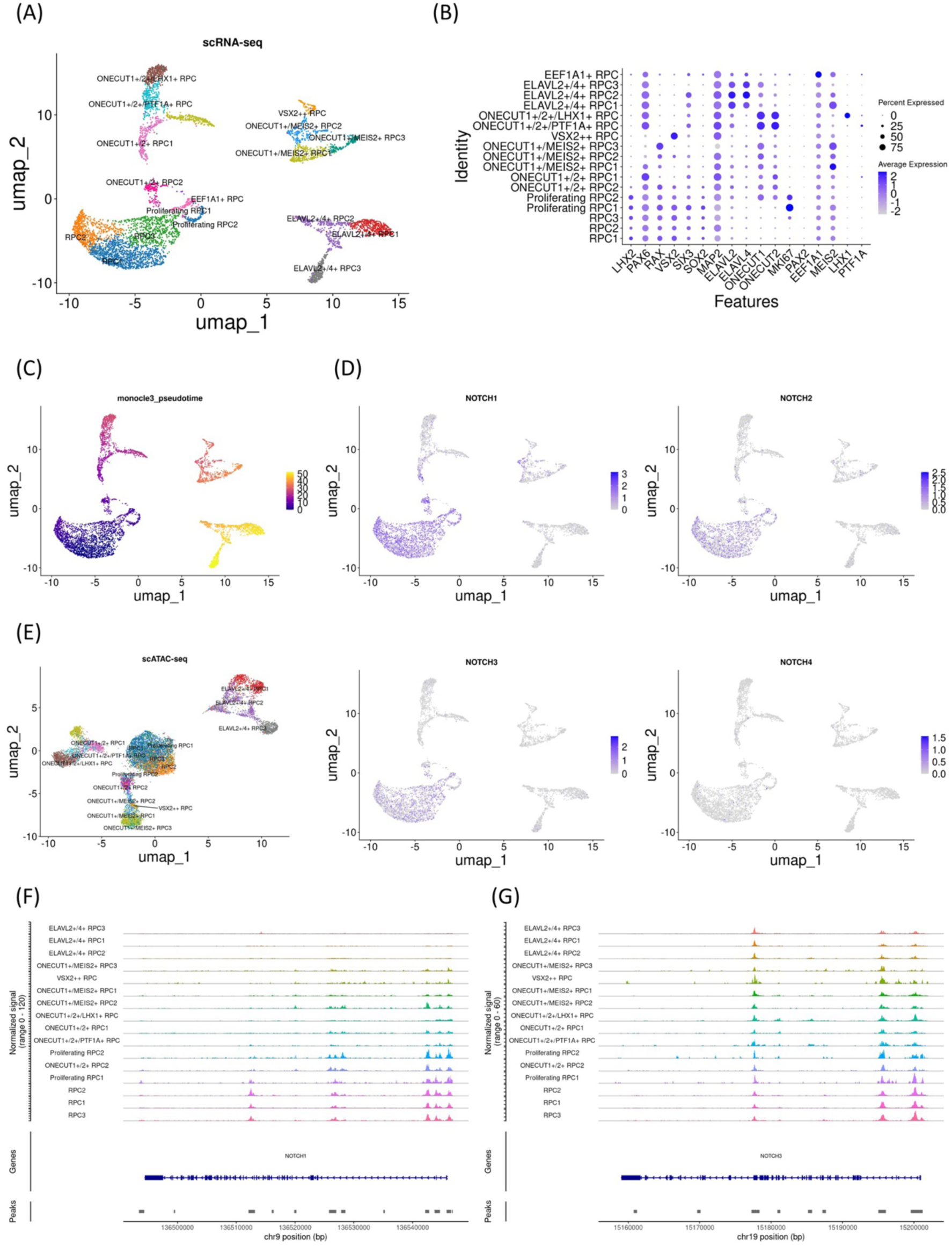
*NOTCH1* mRNA expression decrease and concomitant chromatin remodeling of the day 78 sample. (A) UMAP analysis of the single-cell RNA-seq data identified 17 clusters. (B) Dot plot of marker genes. Note that early RPC clusters expressed *LHX2*, *PAX6*, *RAX*, and *VSX2*. (C) Monocle3 pseudotime analysis. (D) Feature plot of *NOTCH1*-*4*. Note that *NOTCH1* and *NOTCH3* expression was prominent in early RPC clusters. (E) UMAP analysis of the single-cell ATAC-seq data which was integrated with the single-cell RNA-seq data. (F) Coverage plot of the *NOTCH1* locus. (G) Coverage plot of the *NOTCH3* locus.

In the early RPCs (RPC1-3), we observed peaks between Chr9 136510000-136520000 of the *NOTCH1* gene, which diminished in the other RPC clusters (Figure 3F). However, we noted that the chromatin accessibility of the upstream region of the *NOTCH1* locus remained high in *ONECUT*-expressing RPCs. In contrast to these populations, the chromatin accessibility of the *NOTCH1* locus in *ELAVL2/4*-expressing RPCs was very low, which was consistent with the low mRNA expression in these RPC populations. In addition, the chromatin accessibility of the *NOTCH3* locus remained unchanged during RPC differentiation (Figure 3G). In summary, examination of all three independent samples suggested that *NOTCH1* mRNA expression decreased, which was concomitant with chromatin remodeling in Chr9 136510000-136520000 of the *NOTCH1* locus. The feature plots of the marker genes and the dot plots of the *NOTCH* genes from single-cell RNA-seq along with coverage plots of *NOTCH2* and *NOTCH4* from single-cell ATAC-seq are available in “Additional_file_3”

## Discussion

The involvement of Notch signaling in cell fate choices is well documented, including in *Drosophila* neurogenesis [29] and mammalian biliary development [30]. Although the regulation of Notch receptor expression is necessary for these processes, to the best of our knowledge, few studies have used genome-wide investigations of the underlying molecular mechanisms. To examine chromatin remodeling in such regulatory mechanisms, we re-analyzed a single-cell RNA-seq and ATAC-seq dataset from developing retinas in which differentiation trajectories were well characterized. By re-analyzing three independent samples, we observed chromatin remodeling in part of the *NOTCH1* locus, concomitant with changes in its mRNA expression during RPC differentiation.

Transcriptional regulation occurs at many levels, including DNA-binding proteins and miRNAs, as well as chromatin remodeling. Importantly, the chromatin accessibility of the upstream regions of the *NOTCH1* locus was unaffected in *ONECUT*-expressing RPCs. Therefore, the observed mRNA changes might be driven by pathways other than chromatin remodeling. Indeed, *NOTCH3* mRNA expression also diminished during differentiation, although we observed no chromatin remodeling in the *NOTCH3* locus. Because chromatin accessibility in *ELAVL2*/*4*-expressing RPCs was very low, this ensured low mRNA expression of *NOTCH1*.

An ophthalmological study revealed that the epigenetic landscape of cell type-specific enhancers shifted during differentiation of RPCs [31]. For example, in the single-cell ATAC-seq data from embryonic day 14.5 mouse retina, motif enrichment for *Lhx2*, *Rax* and *Pax6* in the early RPCs were observed, and footprinting analysis validated binding of those transcription factors to their motifs. These high chromatin accessibilities decreased as they differentiated into retinal ganglion cells and non-retinal ganglion cells. Although that study is excellent in providing comprehensive and in-depth insights, ours is unique in focusing on the Notch loci for clarifying the regulatory mechanisms in view of Notch signaling biology.

## Limitations

This study elucidated the chromatin regions in which chromatin accessibility differed among RPC subsets. We identified the transition of Notch receptor expression and the accompanying chromatin remodeling of Notch loci in the early stages of RPC differentiation. Further investigations, such as a large deletion of these regions, will be needed to evaluate the contribution of these regions to the differential expression of Notch receptors in RPC subsets.

## Abbreviations

RPC: Retinal progenitor cell

UMAP: Uniform manifold approximation and projection

## Declarations

### Ethics approval and consent to participate

Not applicable.

### Consent for publication

Not applicable.

### Availability of data and materials

A single-cell multiomics dataset (GSE183684) [18] was downloaded from the Gene Expression Omnibus database (https://www.ncbi.nlm.nih.gov/geo/). The code used in the present study is available at our GitHub repository (https://github.com/Yoshitokky/Notch_multiomics_retina/tree/v1.0) that was archived using Zenodo (https://doi.org/10.5281/zenodo.14058572).

### Authors’ contributions

Y. W. performed the analyses and wrote the original manuscript. S.K. assisted in writing the manuscript. T.N. designed the study. S.T. revised and supervised the study. M.Y. designed the study, performed analyses, wrote the original manuscript, and acquired funding.

## Acknowledgements

The authors thank Editage (https://www.editage.jp) for their support with English language editing.

## Supplementary file

File name: Additional_file_1

File format: JPG

Title of data: Supplementary Figure for day 59 data

Description of data: (A) Feature plot of the marker gene mRNA expression. (B) Dot plot showing Notch mRNA expression. (C) Chromatin accessibility at the *NOTCH2* locus as revealed by coverage plots. (D) Coverage plot of the *NOTCH4* locus.

File name: Additional_file_2

File format: JPG

Title of data: Supplementary Figure for day 74 data

Description of data: (A) Feature plot of the marker gene mRNA expression. (B) Dot plot showing Notch mRNA expression. (C) Chromatin accessibility at the *NOTCH2* locus as revealed by coverage plots. (D) Coverage plot of the *NOTCH4* locus.

File name: Additional_file_3

File format: JPG

Title of data: Supplementary Figure for day 78 data

Description of data: (A) Feature plot of the marker gene mRNA expression. (B) Dot plot showing Notch mRNA expression. (C) Chromatin accessibility at the *NOTCH2* locus as revealed by coverage plots. (D) Coverage plot of the *NOTCH4* locus.

