## Supplementary figures and images for "Expression status transition of *NOTCH1* accompanies chromatin remodeling in human early retinal progenitor cells"

### Additional_file_1

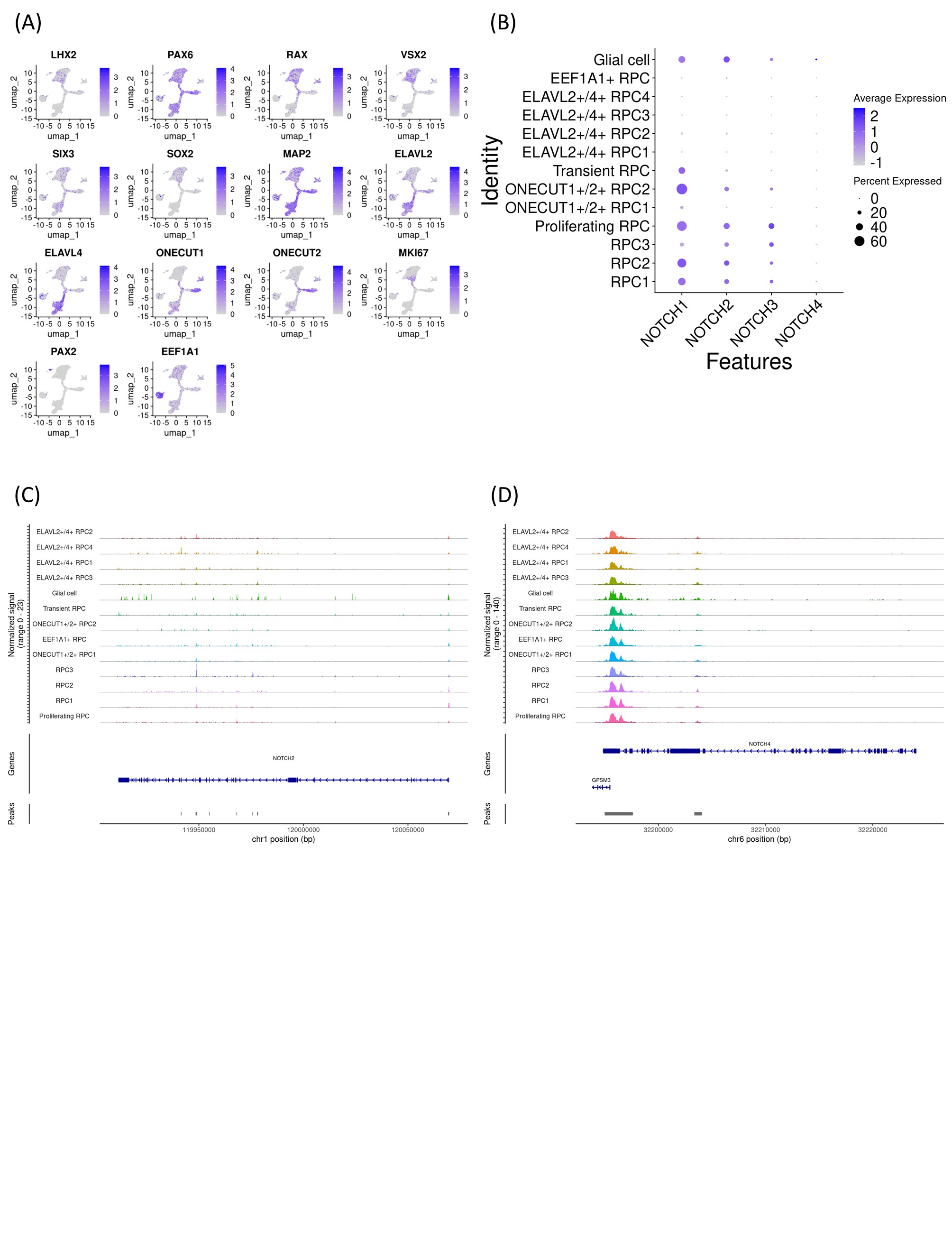

### Additional_file_2

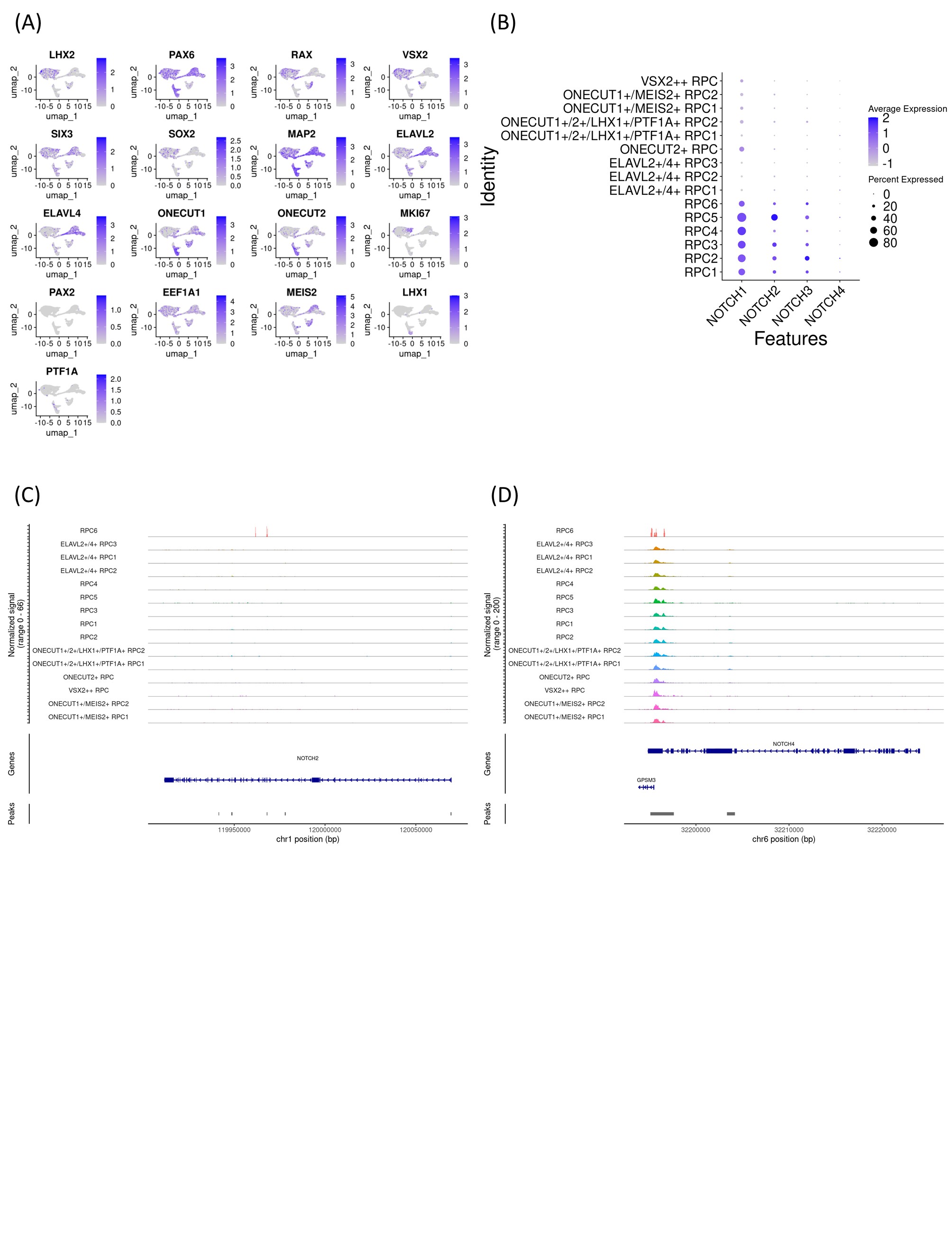

### Additional_file_3

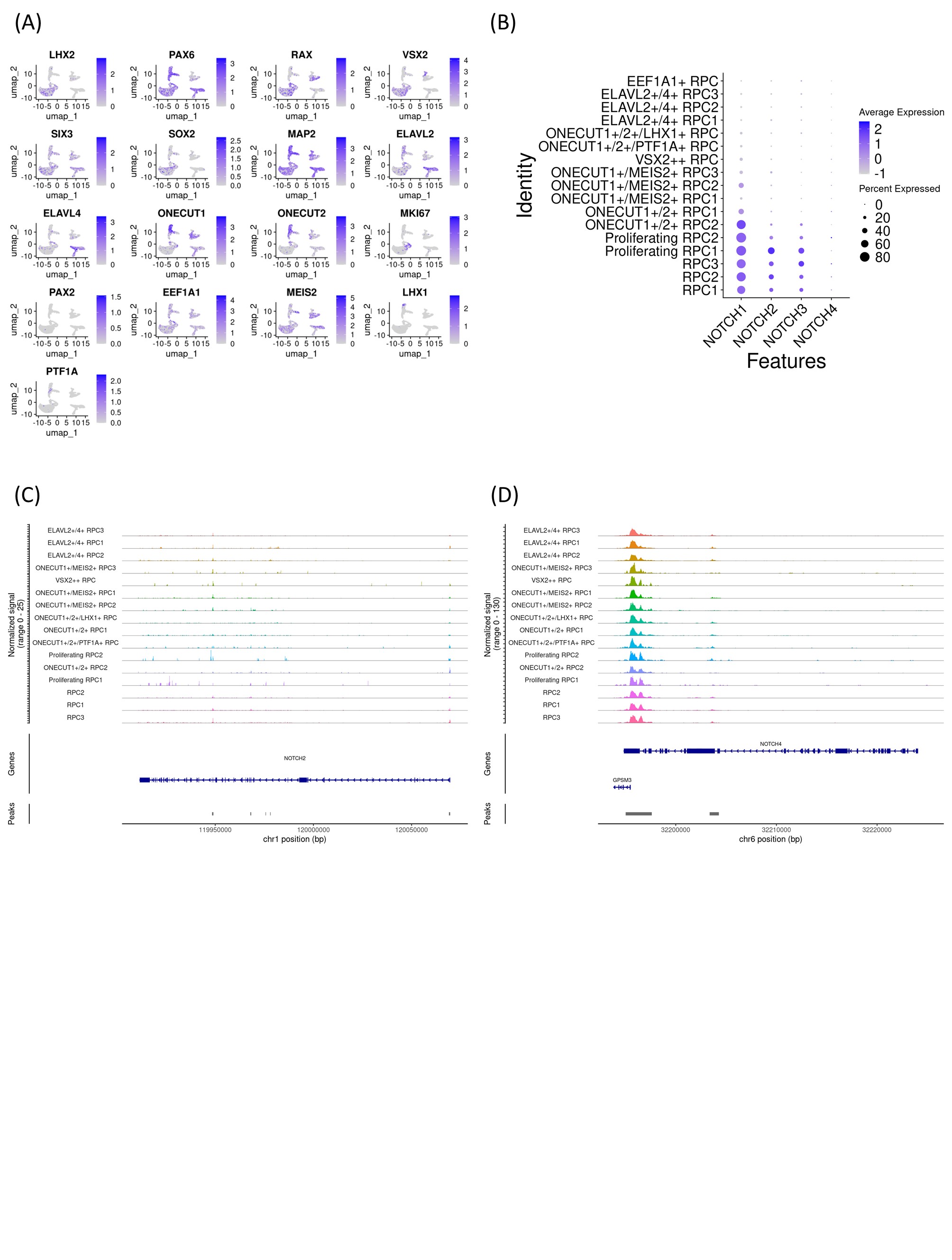
